# Binding of Human ACE2 and RBD of Omicron Enhanced by Unique Interaction Patterns Among SARS-CoV-2 Variants of Concern

**DOI:** 10.1101/2022.01.24.477633

**Authors:** Seonghan Kim, Yi Liu, Matthew Ziarnik, Yiwei Cao, X. Frank Zhang, Wonpil Im

## Abstract

The 2019 coronavirus disease (COVID-19) pandemic has had devastating impacts on our global health. Severe acute respiratory syndrome coronavirus 2 (SARS-CoV-2), the virus causing COVID-19, has continued to mutate and spread worldwide despite global vaccination efforts. In particular, the Omicron variant, first identified in South Africa in late November 2021, has now overtaken the Delta variant and become the dominant strain worldwide. Compared to the original strain identified in Wuhan, Omicron features 50 genetic mutations, with 15 mutations in the receptor-binding domain (RBD) of the spike protein, which binds to the human angiotensin-converting enzyme 2 (ACE2) receptor for viral entry. However, it is not completely understood how these mutations alter the interaction and binding strength between the Omicron RBD and ACE2. In this study, we used a combined steered molecular dynamics (SMD) simulation and experimental microscale thermophoresis (MST) approach to quantify the interaction between Omicron RBD and ACE2. We report that the Omicron brings an enhanced RBD-ACE2 interface through N501Y, Q493K/R, and T478K mutations; the changes further lead to unique interaction patterns, reminiscing the features of previously dominated variants, Alpha (N501Y) and Delta (L452R and T478K). Our MST data confirmed that the Omicron mutations in RBD are associated with a five-fold higher binding affinity to ACE2 compared to the RBD of the original strain. In conclusion, our result could help explain the Omicron variant’s prevalence in human populations, as higher interaction forces or affinity for ACE2 likely promote greater viral binding and internalization, leading to increased infectivity.

**TOC GRAPHIC:** 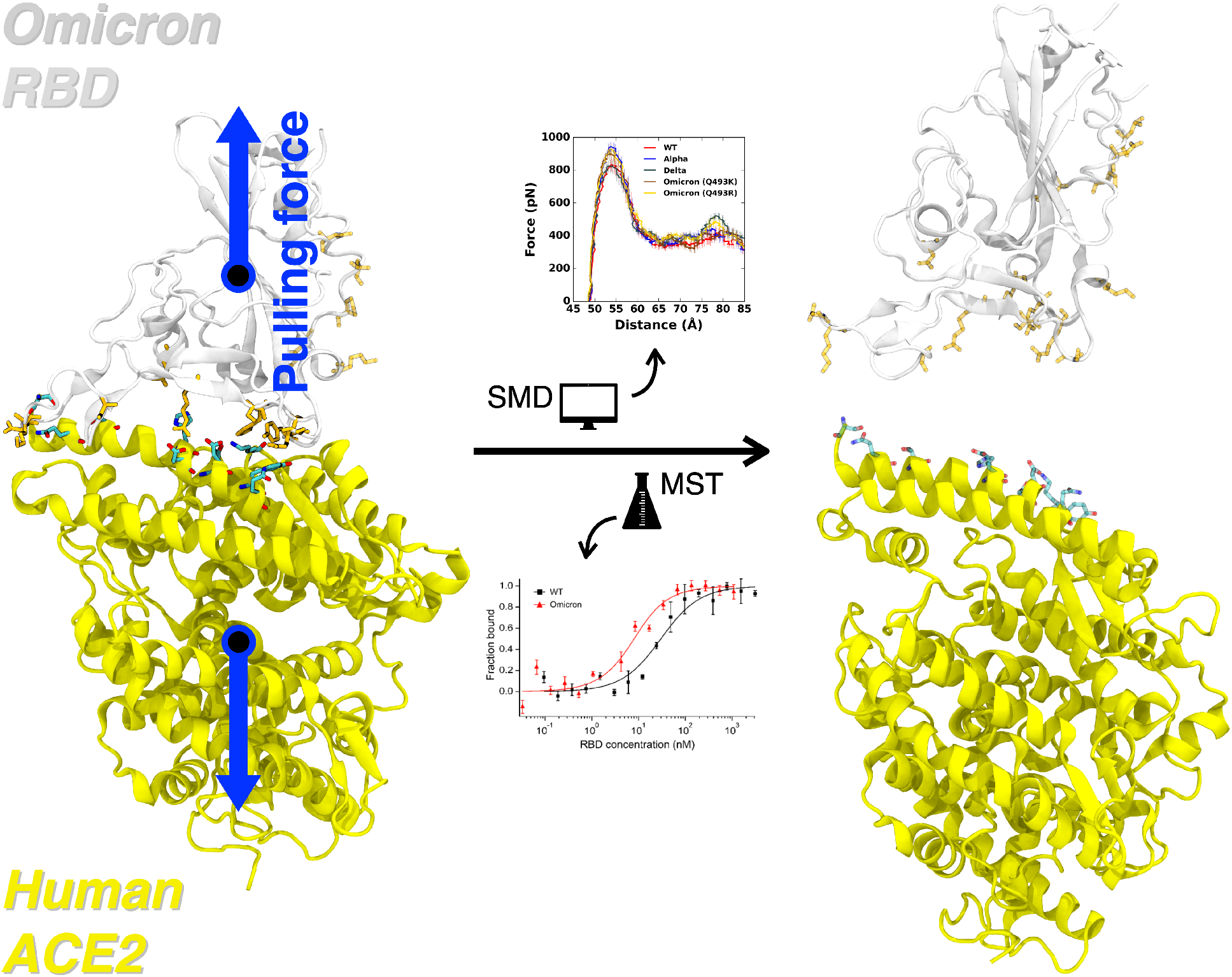

## INTRODUCTION

Severe acute respiratory syndrome coronavirus 2 (SARS-CoV-2) is a positive-sense RNA virus causing the COVID-19 pandemic.^1^ Although vaccines for the virus have been administered to the public, several variants of concern (VOCs) defined by the World Health Organization (WHO), including the Alpha (B.1.1.7), Beta (B.1.351), Gamma (P.1), and Delta (B.1.617.2), as well as the recently identified Omicron variant (B.1.1.529), have caused an increase in infectivity, lowering the efficacy of current vaccines.^2,3^

The Omicron variant was first identified in South Africa in late November 2021. It has overtaken the Delta variant and become the dominant strain circulating in many countries, including theUnited States, causing record-high new COVID-19 cases in December 2021 and January 2022.^3,4^ The Omicron variant features 50 genetic mutations compared to the original (i.e., wild-type, WT) strains identified in Wuhan, China, with 36 mutations located in the spike (S) glycoprotein, over three times more than the number of mutations identified in the four other VOCs.^5^ More specifically, the receptor-binding domain (RBD) of the Omicron spike contains 15 mutations, much more than the 1, 3, 3, and 2 mutations found in RBDs of the Alpha, Beta, Gamma, and Delta variants, respectively.^6^ The RBD is the protein structure used by SARS-CoV-2 to bind to the human angiotensin-converting enzyme 2 (ACE2) receptor to gain host entry.^7^ Such a large number of mutations on the Omicron RBD suggest potential changes in both protein structure and binding affinity to hACE2. Indeed, several recent studies have tackled these issues. Han et al. reported the X-ray crystallography structure and biophysical binding kinetics of the Omicron RBD-ACE2 complex and claimed that Omicron RBD interacts with ACE2 at a similar affinity compared to that of the WT RBD.^8^ However, another protein crystallography and experimental study by Lan et al.^9^ and an MD simulation study by Lupala et al.^10^ reported that mutations in the Omicron RBD resulted in stronger binding toward ACE2. Therefore, the answer to the critical question of whether the Omicron mutations influence the strength of the RBD-ACE2 interaction remains inconclusive.

We recently reported a combined all-atom steered molecular dynamics (SMD) simulation and experimental microscale thermophoresis (MST) approach in studying the binding between human ACE2 and the RBDs of SARS-CoV-2 VOCs Alpha, Beta, Gamma, and Delta in addition to Epsilon and Kappa.^11^ In this study, we applied the same approach to study the Omicron RBD-ACE2 interaction. Our study reveals that the Omicron mutations lead to an enhancement of binding between RBD and ACE2 by forming unique interaction patterns, which are caused mainly by N501Y, Q493K/R, and T478K mutations. This study also provides a better understanding of the role of each mutation in terms of RBD-ACE2 interaction at the molecular level.

## METHODS

### Computational Methods

A fully-glycosylated SARS-CoV-2 RBD and ACE2 complex model was achieved from CHARMM-GUI Archive COVID-19 Protein Library (6vsb_1_1_1_6vw1.pdb).^12^ The model complex includes five and one N-linked glycans in ACE2 (Asn53, Asn90, Asn103, Asn322, and Asn546) and RBD (Asn343), respectively. CHARMM-GUI *Solution Builder*^*13*^ and *Input Generator*^*14*^ were used for system generation. In each variant, the Alpha includes N501Y mutation, Delta has L452R and T478K mutations, and Omicron contains 15 mutated amino acids, i.e., G339D, S371L, S373P, S375F, K417N, N440K, G446S, S477N, T478K, E484A, Q483K/R, G496S, Q498R, N501Y, and Y505H (see also **Figures 1B** and **2D**).^8,15^ Thus, corresponding residues were mutated during the system generation. The CHARMM36(m) force field was used for protein and carbohydrates.^16,17^ The TIP3P water model^18^ was employed with 0.15 M of K^+^ and Cl^-^ ions for mimicking a physiological condition. A large enough system size (190 Å ×190 Å × 190 Å) was considered to make both proteins to be sufficiently solvated when they are fully dissociated. The number of atoms in each system is approximately 550,000.

**Figure 1.**
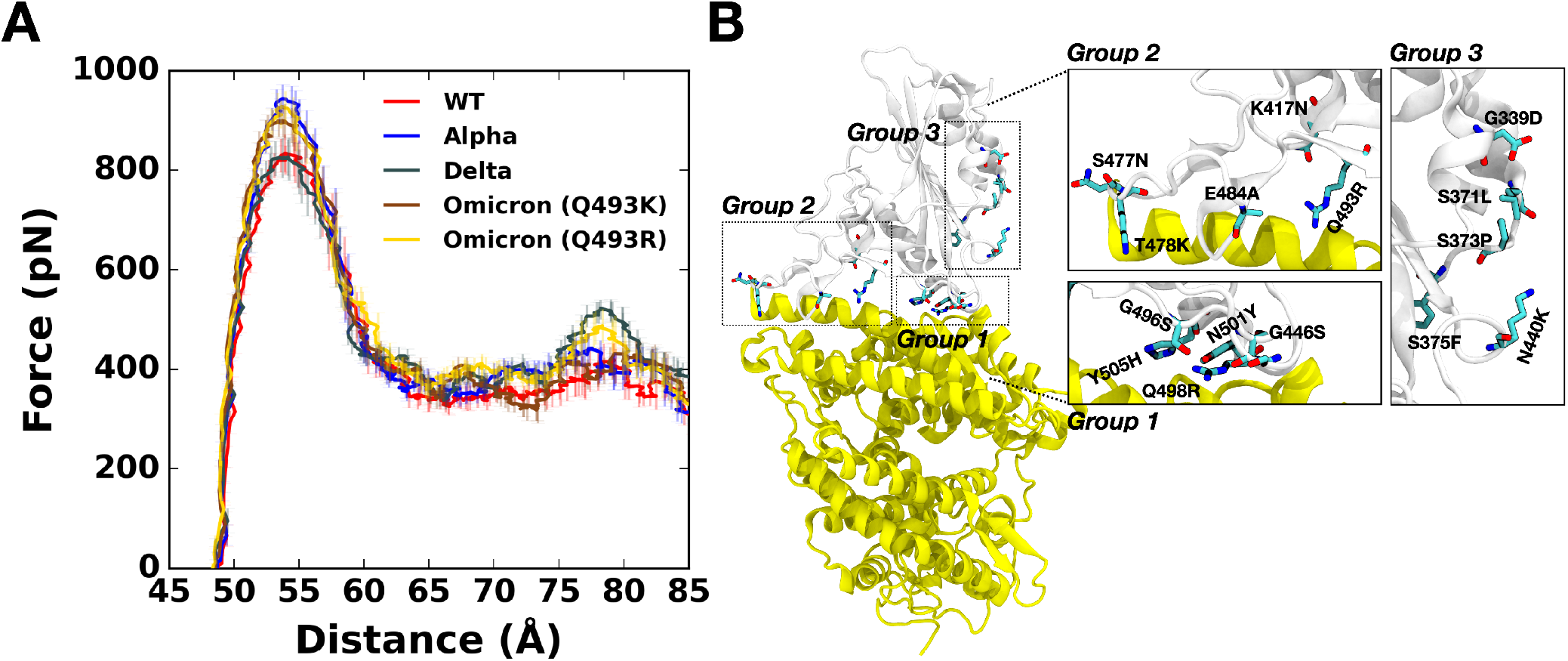
(A) Average force profiles of WT (red), Alpha (blue), Delta (gray), and Omicron variants (brown for Q493K and gold for Q493R) as a function of the distance between the center of mass of ACE2 and RBD. (B) The initial structure of Omicron (Q493R). Mutated residues from the WT are shown as solid sticks that are grouped and magnified into three panels with labels on the right. RBD and ACE2 are respectively colored in light gray (top) and yellow (bottom). All N-glycans, water, and ions are hidden for clarity.

**Figure 2.**
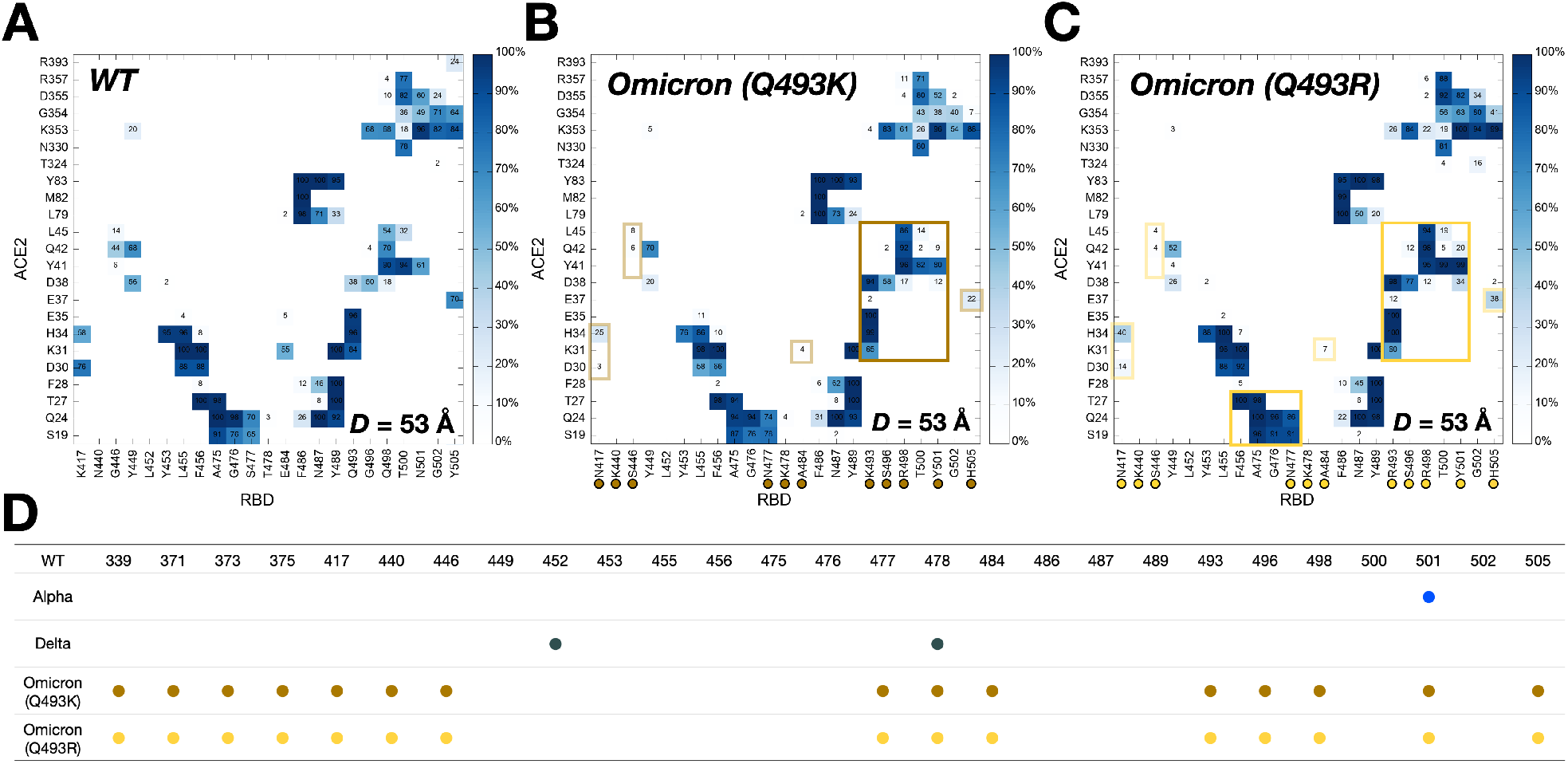
Two-dimensional contact maps of residues in RBD-ACE2 interface at *D* = 53 Å. (A) Interacting residue pairs between RBD^WT^ and ACE2. RBD residues subjected to mutation are marked with colored circles at the bottom: (B) brown for Omicron Q493K and (C) gold for Omicron Q493R. The contact frequency is displayed with colors from light blue to dark blue. Solid and transparent boxes on the map respectively represent increased and decreased interactions between RBD and ACE2 upon mutations. (D) A summary of RBD mutations in the Omicron variants.

The overall simulation details are similar to our previous works.^11,19^ We used NAMD simulation software for the equilibrium and pulling simulations with the COLVARS method.^20^ Initially, the SARS-CoV-2 RBD and ACE2 proteins were aligned and maintained along the X-axis by applying the external forces to the center of mass (COM) of each protein. The effective pulling force applied to the COMs of both proteins was calculated through the following equation:

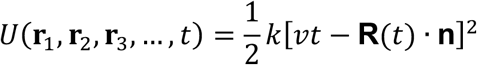

where *k* is the spring constant, *v* is the moving speed of the dummy atoms (i.e., spring potentials), **R**(*t*) is the position of the selected protein COM, and ***n*** is the COM-COM unit vector. This force allows the spring-connected proteins to be pulled in the opposite directions. The moving speed of proteins was set to 0.5 Å/ns, and 5 kcal/mol/Å^2^ of the spring constant was applied to the COM of each protein to let both proteins move along the X direction. To improve statistical results, 20 independent simulation runs of two Omicron variants, Q493K and Q493R, were performed for at least 40 ns to secure that the RBD and ACE2 are entirely dissociated from each other. Note that we used our previous results^11,19^ for WT, Alpha, and Delta for comparison in this study.

The van der Waals interactions were smoothly switched off over 10-12 Å using a force-based switching function^21^ and the particle-mesh Ewald method^22^ with a mesh size of 1 Å was used for the electrostatic interactions. The SHAKE algorithm^23^ was employed to constrain bond lengths including hydrogen atoms. The Langevin piston method and the Langevin damping control method were respectively used for pressure control (1.01325 bar) and simulation temperature (303.15 K).^24^ The NVT ensemble with positional and dihedral restrained was applied for the equilibration simulations, and the NPT ensemble was applied for pulling simulations. The 4 fs simulation time-step was used with the hydrogen mass repartitioning method.^25,26^

### Experimental Methods

Recombinant human ACE2 protein (GenBank accession: AF291820.1, Sino Biological 10108-H08H, Wayne, PA) was labeled with RED-NHS (2nd Generation) dye using the Monolith Protein Labeling Kit (NanoTemper Technologies, MO-L011, München, Germany). Labeled ACE2 (5 nM, final concentration) was mixed with the RBD proteins in a serial 15-step 2-fold dilution starting from 4 μM (for WT) or 1 (for Omicron) μM in PBS buffer supplanted with 0.1 % Pluronic® F-127. Both the WT and Omicron RBD proteins were from ACRObiosystems, Newark, DE (WT: SPD-C52H3, GenBank accession: QHD43416.1; Omicron: SPD-C522e). The mixed RBD+ACE2 samples were separately loaded into 16 premium glass capillaries (NanoTemper Technologies, MO-K025). The 16 capillaries were then placed in the reaction chamber in the order from low to high concentration. MST measurements were conducted on a Monolith NT.115 instrument (NanoTemper Technologies) at 20% excitation power at 24 °C. The measurement was repeated at least three times. K_d_ calculations were performed using the MO Affinity Analysis software (NanoTemper Technologies).

## RESULTS AND DISCUSSION

### The Omicron exhibits stronger RBD-ACE2 binding than WT

Pulling force analysis was performed (**Figure 1A**) as a function of distance, *D*, between COMs of RBD^Omicrons^ and ACE2 to obtain molecular-level insight into the Omicron variants (Q493K/R). Like our previous study,^11,19^ we utilized our fully-glycosylated S RBD-ACE2 complex model for the pulling simulation.^12^ Compared to WT, the Omicron Q493K/R present increased force profiles evidenced by both the first (*D* = 53 Å) and second maximum (*D* = 79 Å) peaks (**Figure 1A**). As shown in **Figure 1B**, there are many mutations in the RBD^Omicron^, and notably, most of the mutations are located at the RBD-ACE2 interface, which can affect the enhanced RBD-ACE2 binding.

### N501Y mutation of the Omicron results in increased maximum force

**Figure 2A,B,C** compare two-dimensional RBD-ACE2 interaction patterns between WT and Omicron variants at *D* = 53 Å. As shown in **Figure 2B,C**, Omicron shows increased contact frequency (solid box) compared to WT (**Figure 2A**), which explains the higher forces than WT at *D* = 53 Å when both proteins are pulled (**Figure 1A**). The contact information is accordant with recent RBD^Omicrons^-ACE2 crystal and cryo-EM structures,^8,9^ indicating that our model system is well validated. To analyze the contact frequency, the number of contacts was calculated between RBD residue 501 (N501 for WT; Y501 for Omicron Q493K/R) and ACE2 in **Figure 3A**. The contact was counted if RBD residue 501 is located within 4.5 Å of heavy atoms of key interacting residues of ACE2. Y501 of Alpha and Omicron variants (Q493K/R) contain more contacts (about 40%) than N501 of WT and Delta. As shown in **Figure 3B,C**, at *D* = 53 Å, Omicron Y501 is positioning closer to ACE2 Y41 and K353 than WT N501, and the π-π and π-cation interactions with neighboring ACE2 Y41 and K353 of the Omicron contribute to hold RBD-ACE2 interface more tightly than WT.

**Figure 3.**
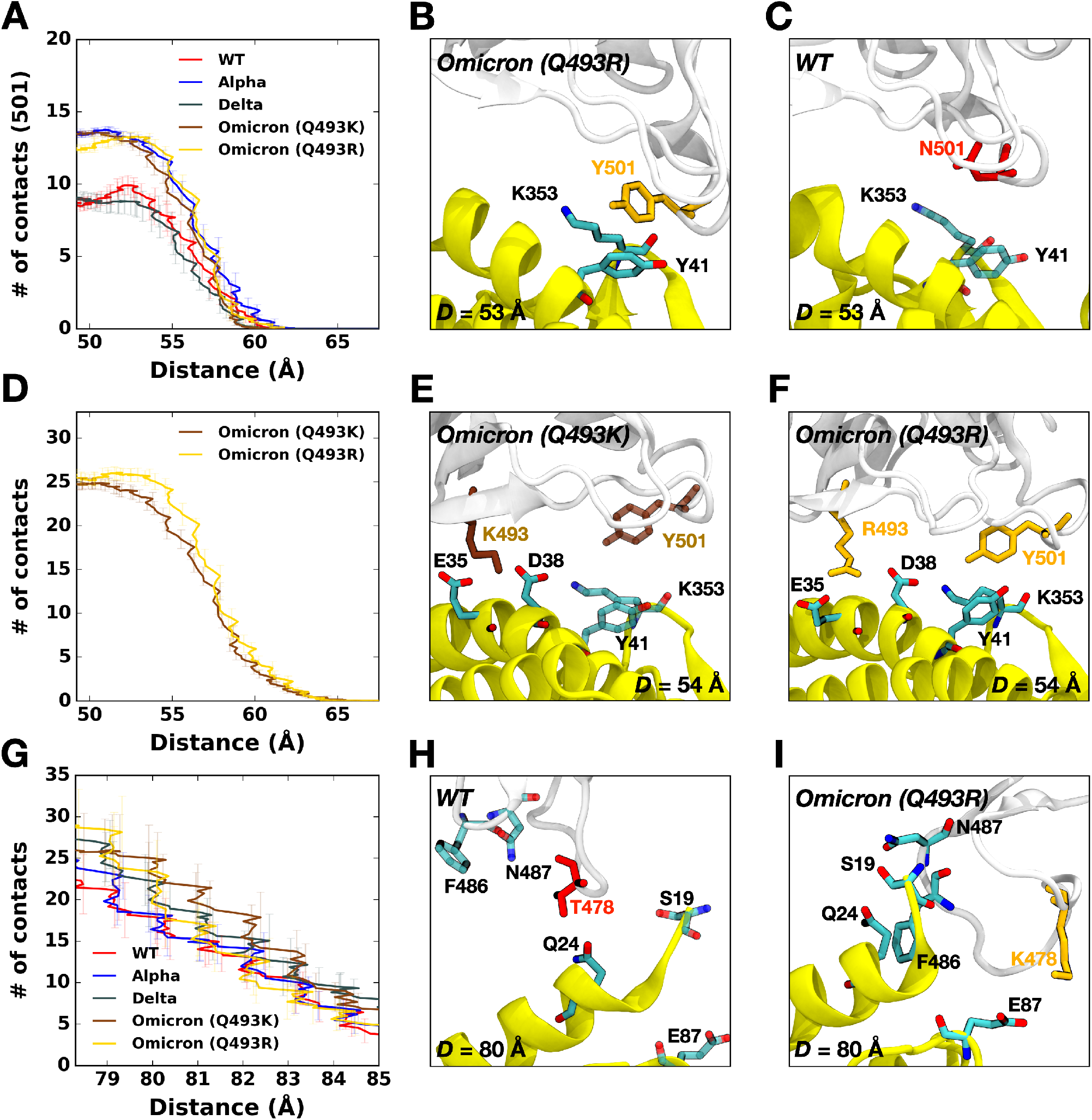
(A) The average number of contacts between RBD residue 501 and ACE2. (B and C) Representative snapshots at *D* = 53 Å of (B) Omicron Q493R and (C) WT. (D) The average number of contacts of Omicron variants between residues in RBD and ACE2 and (E and F) their interacting snapshots of (E) Q493K and (F) Q493R at *D* = 53 Å. (G) The average number of contacts between RBD residues and ACE2 and (H and I) key interaction pairs at *D* = 80 Å of (H) WT and (I) Omicron Q493R. The overall color scheme is the same as in **Figure 1** (i.e., red for WT, brown for Omicron Q493K, and gold for Omicron Q493R). Interacting residues are presented as the solid sticks, and residues losing their interactions are depicted as the transparent sticks.

### The Omicron Q493R mutation also induces reinforced RBD-ACE2 interface

Recent studies report that the Omicron contains either Q493K or Q493R, while other mutations remain the same.^8,15,27-29^ Therefore, we built up two independent Omicron model systems retaining Q493K or Q493R to address both cases (**Figure 2D**). The force profiles of Omicron Q493K at *D* = 53 Å (**Figure 1A**) present weaker maximum forces than Omicron Q493R, although both show higher forces than WT at the same distance. **Figure 2B,C** show the two-dimensional contact maps of Omicron Q493K and Q493R. The overall interaction patterns are similar, but a few differences result in the gap in the force profile. **Figure 3D** shows the number of contact analysis between RBD Omicron Q493K/R and selected residues in ACE2. From the initial state (about *D* = 50 Å), Q493R displays more contacts with ACE2 than Q493K up to *D* = 58 Å. As shown in **Figure 3E**, a nitrogen atom positioning at the end of K493 holds two ACE2 residues, E35 and E38, by forming hydrogen bonds with four oxygen atoms of both E35 and E38. However, R493, which contains two nitrogen atoms, interacts with both D35 and D38 through hydrogen bonds via four oxygen atoms of it, showing increased contact frequency and maintaining RBD Y501 interactions with ACE2 Y41 and K353. This difference induces the gap in the number of contact analysis (**Figure 3D**) and makes RBD^Omicron-Q493R^-ACE2 interface stronger than RBD^Omicron-Q493K^-ACE2 interface. The stronger R493 interaction of Omicron Q493R further brings about increased contacts between RBD (F456, A475, G476, and N477) and ACE2 (S19, Q24, and T27) compared to WT or Omicron Q493K (**Figure 2C**). During the pulling process, the Group 1 in **Figure 1B** is dissociated first, followed by the Group 2 (**Figure S2**), and thus, each group plays a role of hinges between RBD and ACE2 and is responsible for having the first and second maximum forces in the force profile (**Figure 1A**). This indicates that the Omicron Q493R mutation contributes to having a reinforced RBD-ACE2 interface.

### The Omicron presents enhanced second maximum force by T478K mutation

Omicron variants (Q493K/R) have shown unprecedented transmissibility compared to any other SARS-CoV-2 variants,^4,30^ and the force profile in **Figure 1A** confirms that the Omicron Q493K/R, compared to WT, present increased force profiles around *D* = 80 Å as well as *D* = 53 Å. To quantify what brings increased second maximum peak of the Omicron, the number of contacts were calculated (**Figure 3G**), where all residues in both RBD and ACE2 interacting with each other were considered, and the residues located within 4.5 Å were considered to be in contact. As shown in **Figure 3H,I**, the WT T478 shows contacts with ACE2 Q24 without having interactions with ACE2 E87 (**Figure 3H**), but the Omicron K478 makes contacts with ACE2 E87 (**Figure 3I**) in addition to having more RBD^Omicron^-ACE2 interactions (N487-S19 and F486-Q24). Recently published studies claimed that the Omicron may^9,10,27-29^ or may not^8^ show enhanced binding compared to WT. Our results showing enhanced RBD^Omicrons^-ACE2 binding are consistent with the previous experimental and computational studies,^9,10,27-29^ and explain the differential interactions of Omicron Q493K/R, although we only considered single RBD out of trimeric SARS-CoV-2 S protein.

### The Omicron RBD manifests enhanced binding affinity toward ACE2

To validate our simulation results further, we conducted an experimental protein binding assay using MST.^31^ MST has been used for the detection of viral protein-receptor interactions.^32^ We have recently used MST to quantify the interaction between human ACE2 and the S protein RBDs of some VOCs and the WT virus.^11^ Herein, we measured the binding affinity between ACE2 and the Omicron RBD bearing the Q493R mutation (**Figure 4A**). Fitting the MST saturation curve to a first-order 1:1 binding kinetics model yielded a binding affinity of 5.5 ± 1.4 nM, indicating that the RBD^Omicron^-ACE2 interaction has a five-fold higher affinity compared to the 27.5 ± 4.8 nM affinity of the RBD^WT^-ACE2 interaction that we reported recently (**Figure 4B**). Consistent with our simulation results, the RBD^Omicron^-ACE2 interaction also has a higher affinity than all the other VOCs that we measured earlier, including the Alpha, Beta, and Delta VOCs (**Figure 4B**).

**Figure 4.**
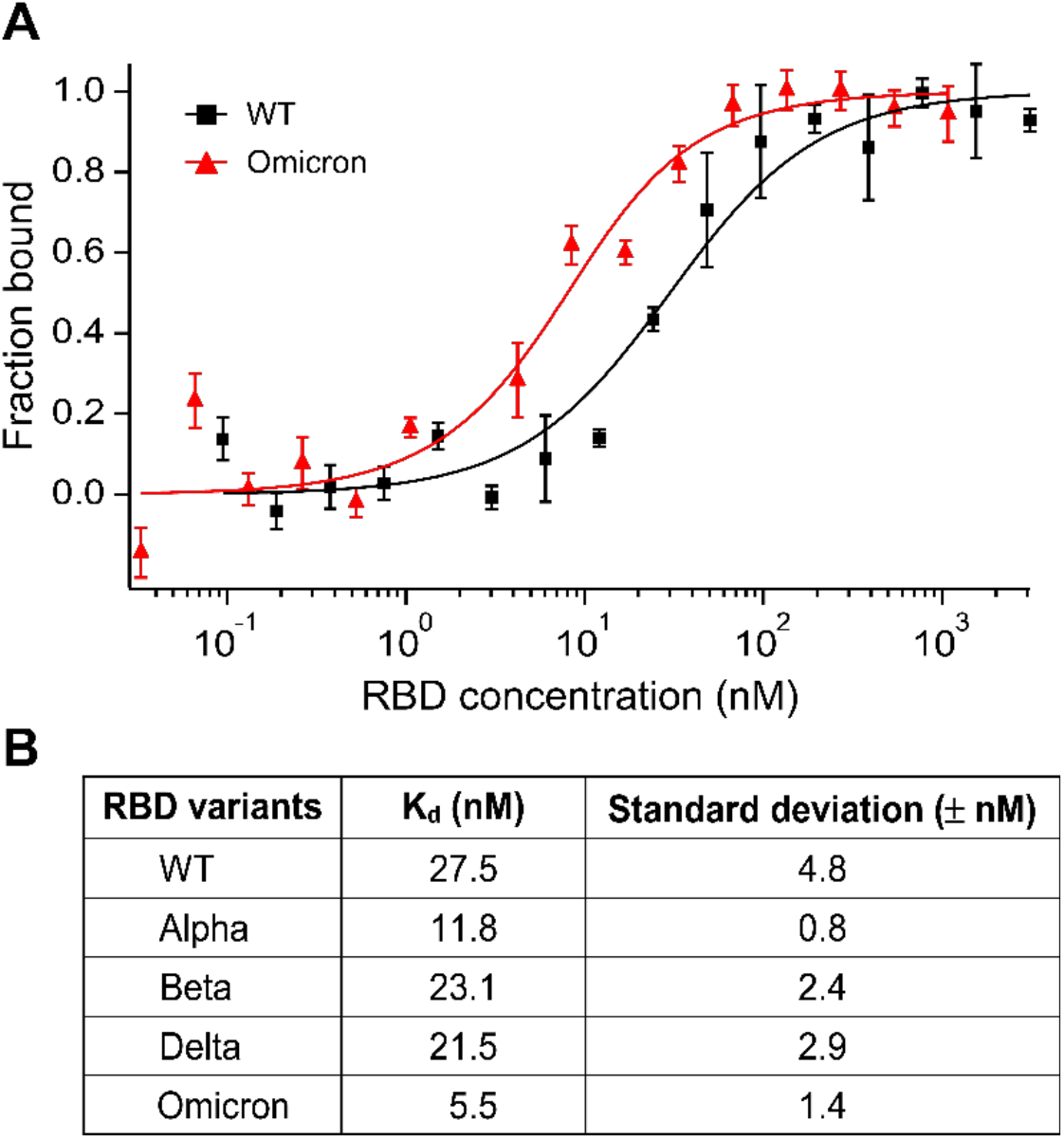
(A) Microscale thermophoresis (MST) analysis of the interaction between ACE2 and RBD of the WT or Omicron variants. Error bars represent standard deviations from five individual repeat measurements. The binding affinities were determined by fitting the data with the ‘K_d_’ model of the MO Affinity software. (B) Affinities of ACE2 binding to RBD variants detected by MST. The MST responses were fitted to the 1:1 binding model. The K_d_ rates are shown as fit ± one standard deviation. Data of the WT, Alpha, Beta, and Delta variants were adapted from our prior study.^11^

## CONCLUSIONS

This study characterizes interactions between ACE2 and the Omicron RBDs (Q493K or Q493R). Both SMD simulation and MST experiment confirm that the Omicron variant RBD displays stronger binding toward ACE2 than the WT RBD. Our analysis shows that the Omicron variants exhibit unique interaction patterns reminiscing the features of previously dominated Alpha (N501Y) and Delta (L452R and T478K) variants. Compared to the WT virus, the N501Y, Q493K/R, and T478K mutations are responsible for the higher force for RBD^Omicrons^-ACE2 dissociation. We hope that this study provides valuable information on the mechanism behind the significantly increased transmissibility of heavily mutated Omicron variants.

## ASSOCIATED CONTENT

The supporting information is available free of charge at https://pubs.acs.org/doi/*.*/acs.jctc*.

Force profiles of all replicas of all variants; two-dimensional contact map of Gamma, Kappa, and Delta; N90-glycan contact; β-strand interactions of WT and Epsilon; MST analysis of variants and affinities.

## ACKNOWLEDGEMENTS

This work was supported in part by NIH AI133634 and NSF 1804117 (to X.F.Z.), NIH R21 AI163708 (to X.F.Z. and W.I.), NIH GM138472, and MCB-1810695 (to W.I.), and an internal grant from Lehigh University (to X.F.Z, and W.I.).

