## Supporting Information for "Binding of Human ACE2 and RBD of Omicron Enhanced by Unique Interaction Patterns Among SARS-CoV-2 Variants of Concern"

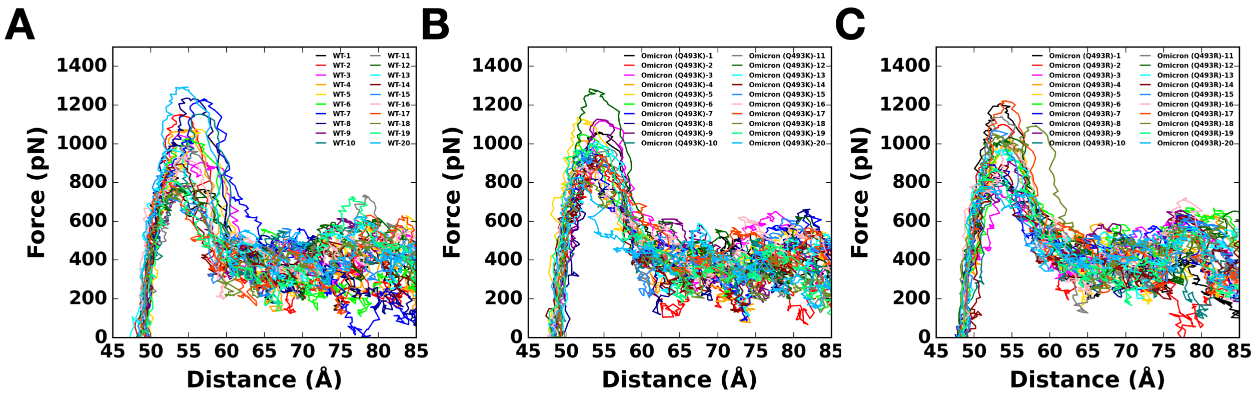


**Figure S1.** Force profiles of 20 replicas of (A) WT, (B) Omicron Q493K, and (C) Omicron Q493R as a function of the distance between the COM of RBD and ACE2.

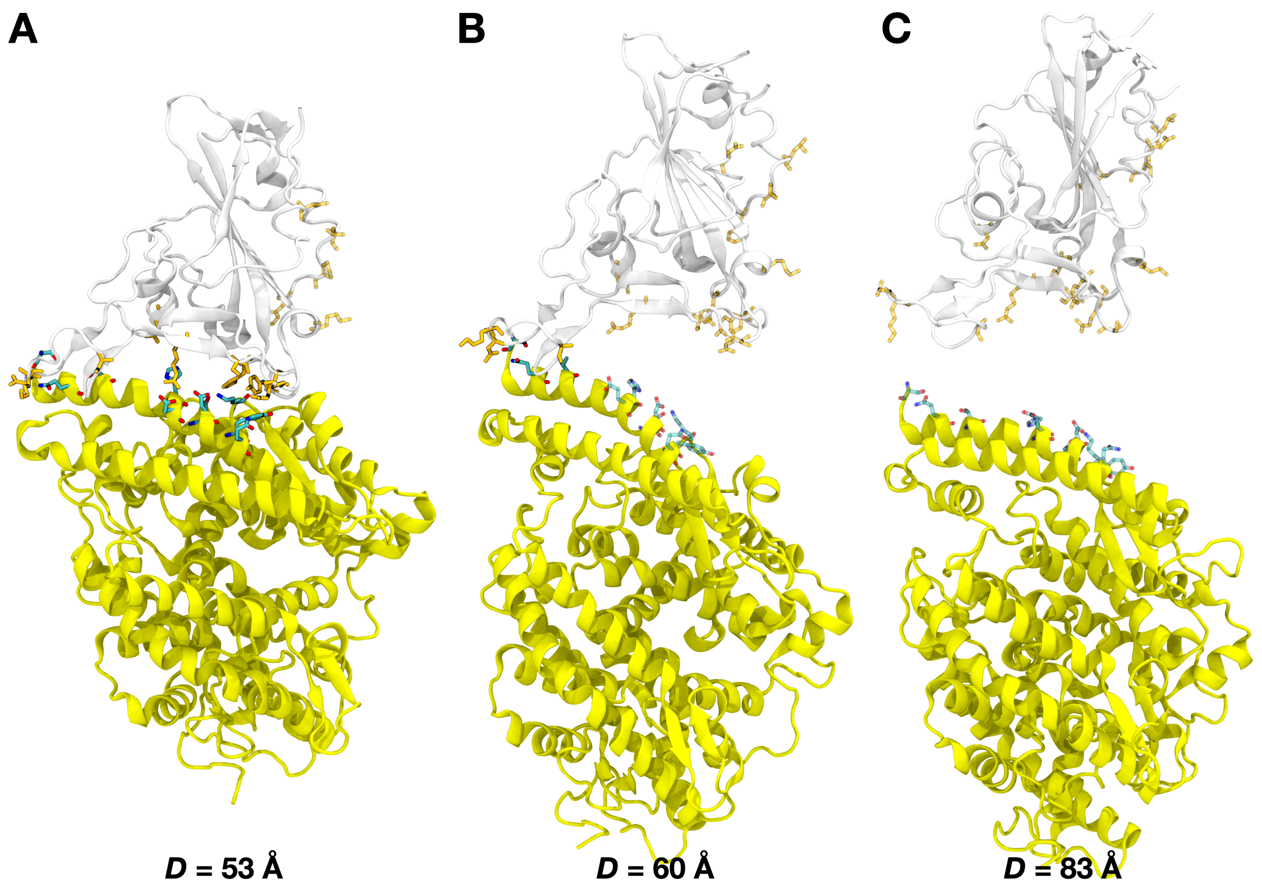


**Figure S2.** Separation process between Omicron RBD and ACE2 at *D* = 53 Å, 60 Å, and 83 Å, respectively. The color scheme is the same as in **Figure 1B**.
